# Scavenger receptor B1 facilitates the endocytosis of *Escherichia coli* via TLR4 signaling in mammary gland infection

**DOI:** 10.1101/2022.09.05.506597

**Authors:** Qamar Taban, Syed Mudasir Ahmad, Peerzada Tajamul Mumtaz, Basharat Bhat, Ehtishamul Haq, Suhail Magray, Sahar Saleem, Nadeem Shabir, Amatul Muhee, Zahid Amin Kashoo, Mahrukh Hameed Zargar, Abrar A. Malik, Nazir A. Ganai, Riaz A. Shah

**Author notes:** **Correspondence Syed Mudasir Ahmad**, Associate Professor, Division of Animal Biotechnology, Sher-e-Kashmir University of Agricultural Sciences and Technology of Kashmir, FV.Sc and A.H, Shuhama, Jammu and Kashmir, India.

## Abstract

SCARB1 belongs to class B of Scavenger receptors (SRs) that are known to be involved in binding and endocytosis of various pathogens. SRs have emerging role in regulating innate immunity and host-pathogen interactions by acting in co-ordination with Toll-like receptors. Little is known about the function of SCARB1 in milk-derived mammary epithelial cells (MECs). This study reports the role of SCARB1 in infection and its potential association in TLR4 signaling on bacterial challenge in Goat mammary epithelial cells (GMECs). The novelty in the establishment of MEC culture lies in the method that aims to enhance the viability of the cells with intact characteristics upto a higher passage number. We represent MEC culture to be used as a potential infection model for deeper understanding of animal physiology especially around the mammary gland. On *E*.*coli* challenge the expression of SCARB1 was significant in induced GMECs at 6 h. Endoribonuclease-esiRNA based silencing of SCARB1 affects the expression of TLR4 and its pathways i.e. MyD88 and TRIF pathways on infection. Knockdown also affected the endocytosis of *E*.*coli* in GMECs demonstrating that *E*.*coli* uses SCARB1 function to gain entry in cells. Furthermore, we predict 3 unique protein structures of uncharacterized SCARB1 (*Capra hircus)* protein. Overall, we highlight SCARB1 as a main participant in host defence and its function in antibacterial advances to check mammary gland infections.

## Introduction

Scavenger receptor B type 1 (SCARB1/SR-BI/CLA1) has been demonstrated to interact with non-self ligands (1, 2, 3, 4, 5) and in particular is known to mediate inflammatory responses and immunomodulatory effects (6). SCARB1 is known to triggers pro-inflammatory signaling to defend against variety of pathogenic microorganisms (7, 8, 9, 10, 11) and microbial components such as lipopolysaccharide (LPS) (12, 13) and has emerged as a potential receptor for attachment and entry for hepatitis C virus (HCV) (14, 15, 16) and severe acute respiratory syndrome coronavirus 2 (SARS-CoV-2) as well (17). Primarily, SCARB1 receptor is known to facilitate the uptake of cholesterol both as free cholesterol (FC) and cholesteryl esters from (Low-density lipoprotein) LDL and (High-density lipoprotein) HDL in mammary epithelial cells (18). But little is known about the immune function in MECs, which act as the first line of defence to guard mammary tissue from exogenous threats and in launching the innate immune response (19, 20). Goat is a model species to study the mammary host-pathogen interactions and is a better choice for modelling the human mammary gland because of the morphological similarities (21). The establishment of a primary mammary epithelial cell (PMEC) line, as a model represents the *in vivo* condition. Our in-house protocol provides a novel non-invasive method to isolate MECs from goat milk with maintainace of the primary culture till a higher passage number i.e. 15 with enhanced viability.

*E. coli*, especially is known to develop severe clinical symptoms in the mammary gland. The role of SCARB1 in the entry of *E. coli* in MECs during infection is significant as bacterial intramammary infections in cattle as well as in humans are a major concern (22, 23). It has led to the highest economic losses in the developing countries and affected the rural economy for several decades (24) So, to understand the host-pathogen signaling mechanisms is necessary to expedite the improvement of approaches at controlling mammary gland infections.

SRs are documented as pattern recognition receptors (PRRs) and known to act in coordination with other PRRs such as TLRs (Toll-like receptors) in generating the immune responses on the challenge with LPS of *E*.*coli* (25, 26). In response to LPS, bacteria-induced inflammation is partially SR-BI/II-dependent with Toll-like receptor 4 (TLR4) in contributing to cytokine secretion (8). So, the focus was on the association of SCARB1 receptor with TLR4 signalling pathways in infection `in dairy ruminants, which has not been highlighted` earlier as most studies have focused on the role of SCARB1 as a lipoprotein receptor. TLR4 is one of the best-characterized TLRs and is mainly activated by the LPS of *E*.*coli* (34). In response to *E. coli* or LPS, BMECs (Bovine mammary epithelial cells) initiate a pro-inflammatory response via nuclear factor kappa B (NF-κB) primarily through TLR4 (27, 28) which follows the elimination of the bacteria (29). On activation, TLR4 acts as a “bipartite receptor” and recruits two adaptor molecules; myeloid differentiation primary response protein 88 (MyD88) (30, 31) and TIR-domain-containing adapter-inducing interferon-β (TRIF) that trigger MyD88 dependent and independent pathways respectively and leads to the secretion of pro inflammatory molecules (32). MyD88-mediated signaling occurs mainly at the plasma membrane (33), which is diverged into NF-*κ*B (41, 44) and the mitogen-activated protein kinase (MAPK) branches via TAK1 (34).

On activation, TLR4 use TRIF to activate an alternative pathway (35) that begins in early endosomes after endocytosis of the receptor (36). LPS is internalized from the plasma membrane into early endosomes along with TLR4 containing phagosomes (37). This follows interferon regulatory factor 3 (IRF3) and (Interferon-β) INF-β production. Since, TIRAP–MyD88 and TRAM–TRIF signal transduction pathways originate at spatially separated cellular locations and sequentially in time. So, we assumed, that internalization/endocytosis of bacteria might be TLR4-TRIF pathway mediated which, is unexplored in MECs.

The primary goal of this study was to test the hypothesis that SCARB1 may be involved in an *E*.*coli*-induced inflammatory response via TLR4 signaling pathways and the potential role of SCARB1 in *E. coli* internalization/endocytosis in GMECs.

## Results

### Establishment of *in vitro* mastitic model of GMECs

#### Establishment of a mammary epithelial cell line from goat milk

The primary culture of GMECs was established from exfoliated MECs of milk and sub-cultured up to passage 15 (P15). Initially, many milk artefacts dominated the flask that was removed during the washing step. A heterogeneous population of cells i.e. fibroblasts were not found at P0 or late passages. Few single epithelial cells were observed at P0 that grew into small epithelial patches after 5 days of culture. After about 10 days of culturing the cells in flasks, a highly confluent flask with monolayer, cobblestone, and epithelial-like morphology was formed. Alveoli structures were also observed which are typical of the mammary epithelial cells **(Fig. 1a, d)**.

**Fig. 1.**
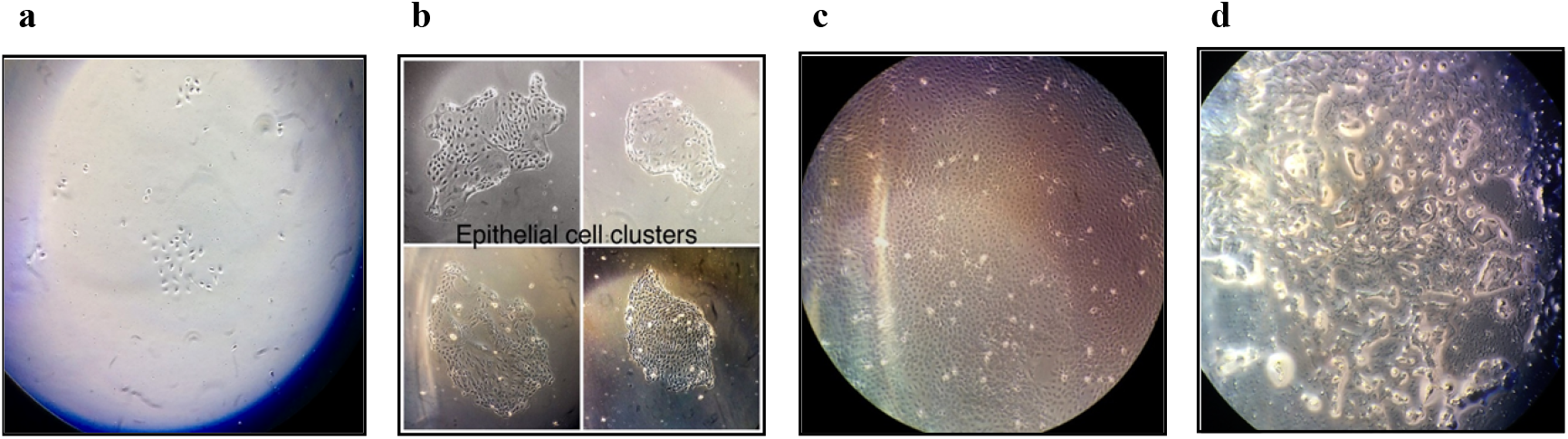
GMECs represent characteristic morphologies of mammary epithelial cells with prominent cell nucleus and the nucleoli. **a**, GMECs on day 3-post seeding (×4). **b**, GMECs form cobblestone like morphology on day 5-post seeding (×4). **c**, Highly confluent GMECs on Day 10 (×4). **d**, Milk droplet like inclusions are observed (×10).

#### Biological characteristics and proliferation rate of GMECs

Early and late passage GMECs showed rapid growth and proliferation during 10 days post-seeding. Trypan blue-based proliferation assay performed on GMECs at passages 2 and 15 showed an “S” shaped growth curve that represents the good ability of cell survival and adherence **(S1, Fig. 1a)**.

MTT assay was used as an index of cell proliferation and viability of GMECs (**S1, Fig. 1b)**. The graph infers absorbance at 570 nm between passages over a span of 7 days for GMEC culture. Quantitative analysis results show linearity in the rate of proliferation based on the metabolic activity inside viable cells.

The nuclear and cytoplasmic morphology of GMECs at various passages was observed after Giemsa staining of cells and that the derived primary cells cultures had no heterogeneous population. GMECs appeared round-shaped, densely packed islands with multiple nucleoli and exhibited typical cobblestone morphology **(S1, Fig. 1c)**.

#### GMECs express epithelial cells specific marker protein CKT-18

The epithelial characteristics of established GMECs were examined by IF for CKT-18 at early passages P2 and P3 and late passages, P15. A monolayer of cells was stained with specific primary antibodies followed by secondary antibodies. The results of immunostaining show that the established cultures are positive for CKT-18 thus confirming their epithelial nature **(Fig. 2a)**. The culture obtained is pure with no heterogeneous population of fibroblasts. A negative control experiment with a specific CKT-18 antibody revealed no positive fluorescence signal for goat tissue fibroblasts **(Fig. 2b)**. While a positive control experiment on MCF-10A revealed a positive fluorescence signal same as that of GMECs **(Fig. 2c)**. However, poor staining of some GMECs might be due to low levels of expression of CKT-18.

**Fig. 2.**
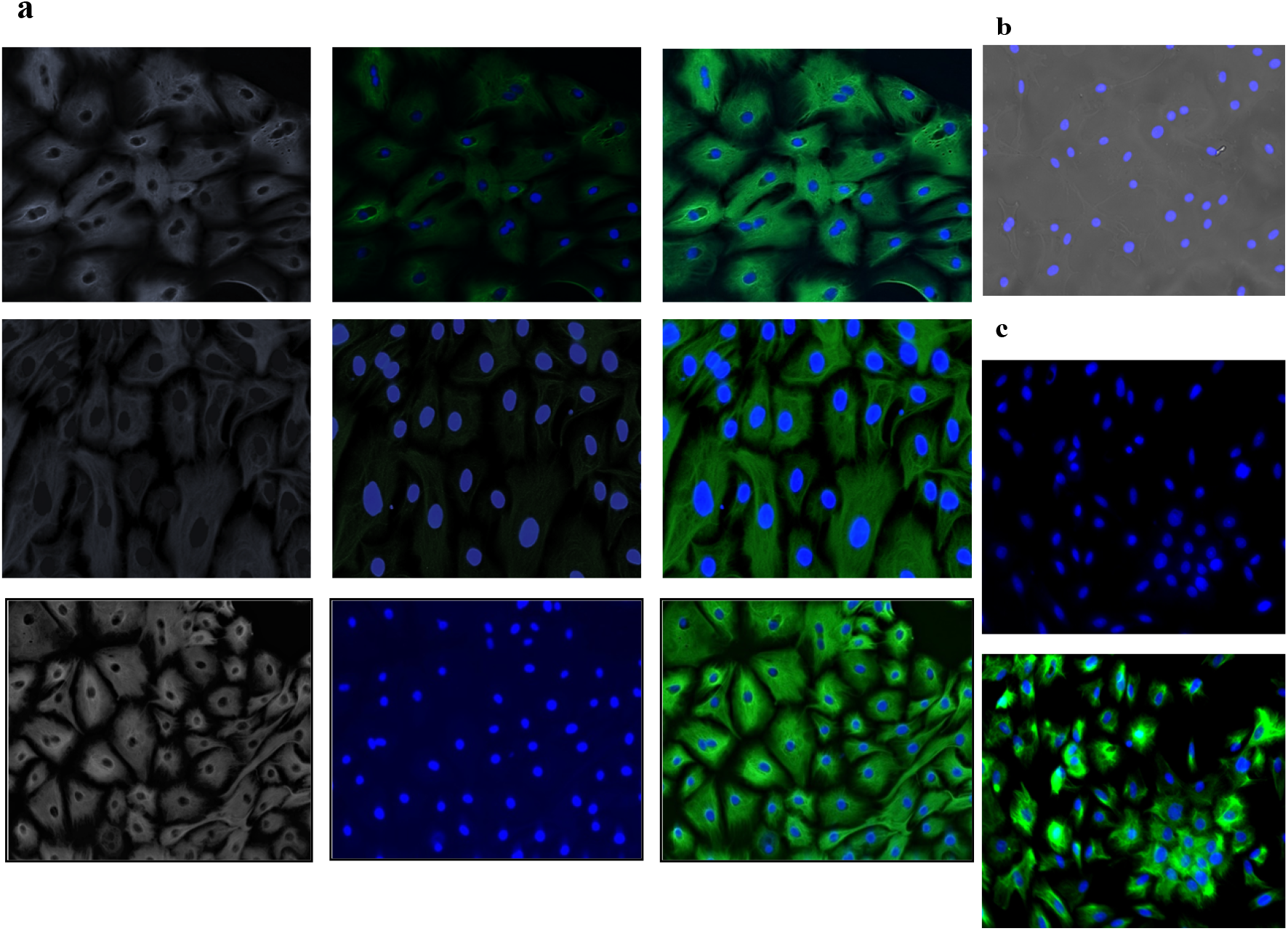
The established GMECs were characterized by immunostaining for epithelial marker protein CKT-18. **a**, Fluorescent images of GMECs at passages P2, P3 and P15 show cells immunostained for CKT-18 with anti-cytokeratin 18 monoclonal antibody *(green)* and nuclei counterstained with DAPI *(blue)*. **b**, Fibroblasts show no immunostaining with specific antibodies for CKT-18. **c**, MCF10A at passage 10 was used as the positive control. Scale 100 μm.

#### Chromosome Analysis

Chromosome analysis of GMECs revealed normal diploid (2n=30) chromosome number specific for *Capra hircus* (66). Representative images of metaphase spread and karyotype are shown in **(S1, Fig. 2)**. Also, there was no instance of chromosomal drift during the course of subculturing to later passages.

#### Differentiation of GMECs

Signs of lactogenic differentiation of mammary cell culture is an expression of milk proteins and changes in MEC morphology. This determines that the cells represent *in vivo* conditions. Expression of the *Capra hircus* β-casein gene (CSN-2) (the most abundant protein in goat milk) was determined in the differentiated MECs by RT-qPCR. Expression of CSN-2 was induced synergistically (1-3.55 fold, *P* < 0.05) by a combination of lactogenic hormones and local growth factors after 5 days **(Fig. 3)**. Lumen-like milk drop–like structures were observed in differentiated cells. Identical morphology of early P2 and late P13 passages after induction was observed **(S1, Fig. 3)**.

**Fig. 3.**
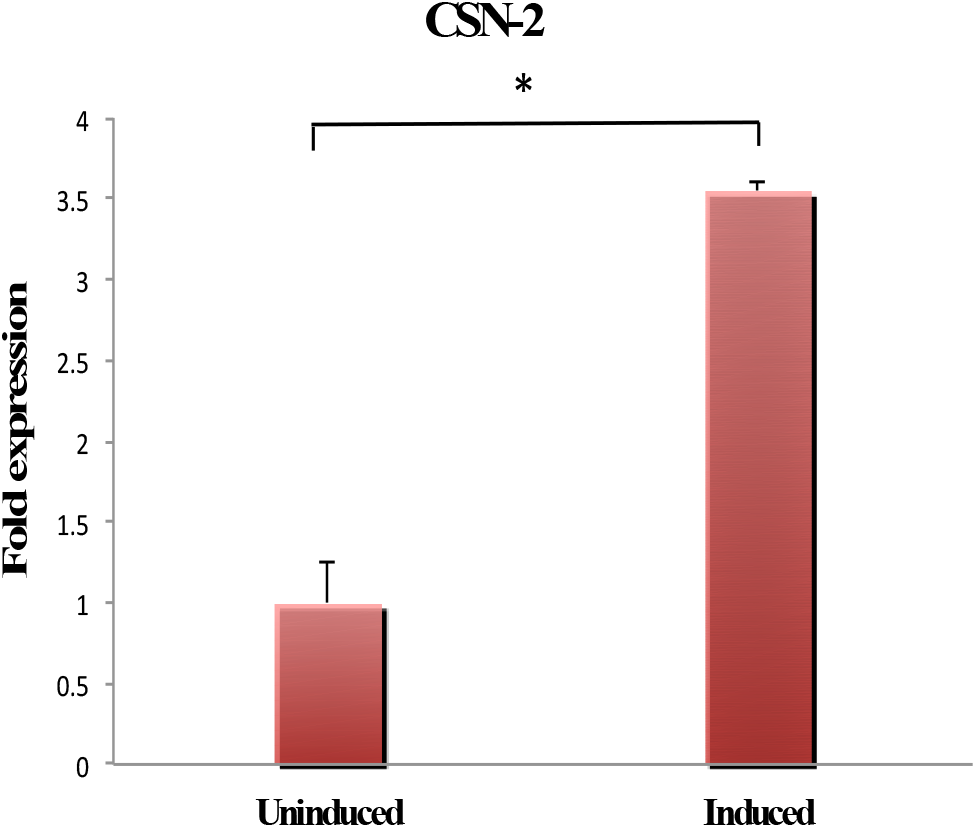
CSN-2 mRNA is highly expressed in differentiated GMECs and RT-qPCR analysis shows a significant increase in the expression of CSN-2 in differentiated GMECs than non-differentiated cells (3.55 fold). Quantitative PCR data were normalized to GAPDH and β-Actin. The values are the mean ± SEM for triplicate samples. \**P* < 0.05.

#### Expression of GFP in GMECs

Green fluorescent protein (GFP) positive GMECs transfected with plasmid EGFPC1 and after 24h, the cells were detected under the Inverted fluorescent microscope. The cells that express GFP appear green in colour **(S1, Fig. 4)**.

#### GMECs response to bacterial challenge

Adhered bacteria can be seen on the surface of giemsa stained cells infected with hk *E*.*coli*. This standard visualization technique gives a qualitative view of adhesion and helps us to distinguish different patterns of adhesion. On giemsa stained cells treated with infection media without bacteria, no adhered bacteria can be seen **(Fig. 4a)**. To confirm the cellular response of bacterial infection, expression of TNF-a and IL8 was assessed through RT-qPCR after 3 h of bacterial challenge with hk *E*.*coli* at 1:100 MOI (Fig. 21). Both the immunogenes showed elevated expression after challenge, thereby validating the cell response after bacterial challenge. Increase in the mRNA expression of TNF-a (35.58 fold, *P* < 0.05 value) **(Fig. 4b)** and IL8 (77.52 fold, *P* < 0.05) **(Fig. 4c)** was seen in GMECs on infection with *E. coli* against non-infected control cells.

**Fig. 4.**
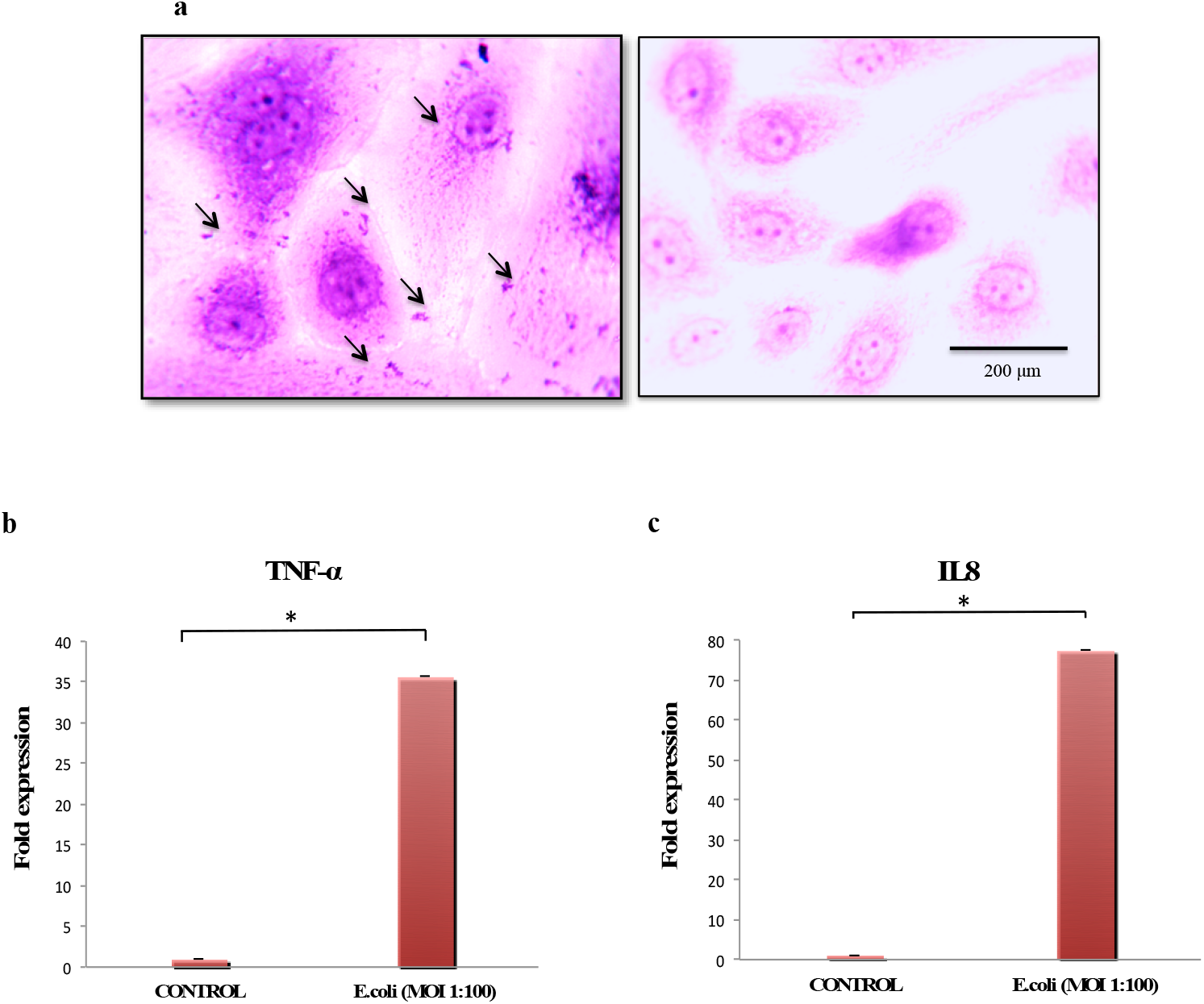
**a**, Adhered *E. coli* on Giemsa stained GMECs. Arrows show the attached bacteria to the cells. A non-infected control was also included (×100). **b**, Fold Expression change of TNF-a (35.58 fold) and IL8 (77.52 fold) genes in *E. coli* challenged GMECs against non-infected control. The y-axis represents the log2 fold change of gene expression. Quantitative PCR data were normalized to GAPDH and β-Actin. The values are the mean ± SEM for triplicate samples. \**P* < 0.05.

### SCARB1 expression

#### In vivo SCARB1 expression

To understand the importance of the SCARB1 receptor in recognizing pathogens and activating downstream signaling; we first detected the expression status of SCARB1 in mammary gland tissue of healthy animals and animals with mammary gland infection/mastitis. To confirm coliform mastitis, milk samples were analyzed, and the *E*.*coli* was identified before tissue collection **(S2, Fig 1)**. After mammary gland tissue collection of mammary gland infected cases, changes in SCARB1 protein levels were detected by western blotting. The SCARB1 protein levels were significantly higher in *E. coli*-mastitic cases compared with healthy goat controls (*P* < 0.05) **(Fig. 5)**. The results indicate that coliform mastitis triggers SCARB1 expression *in vivo* and this receptor is involved in infection in dairy goats.

**Fig. 5.**
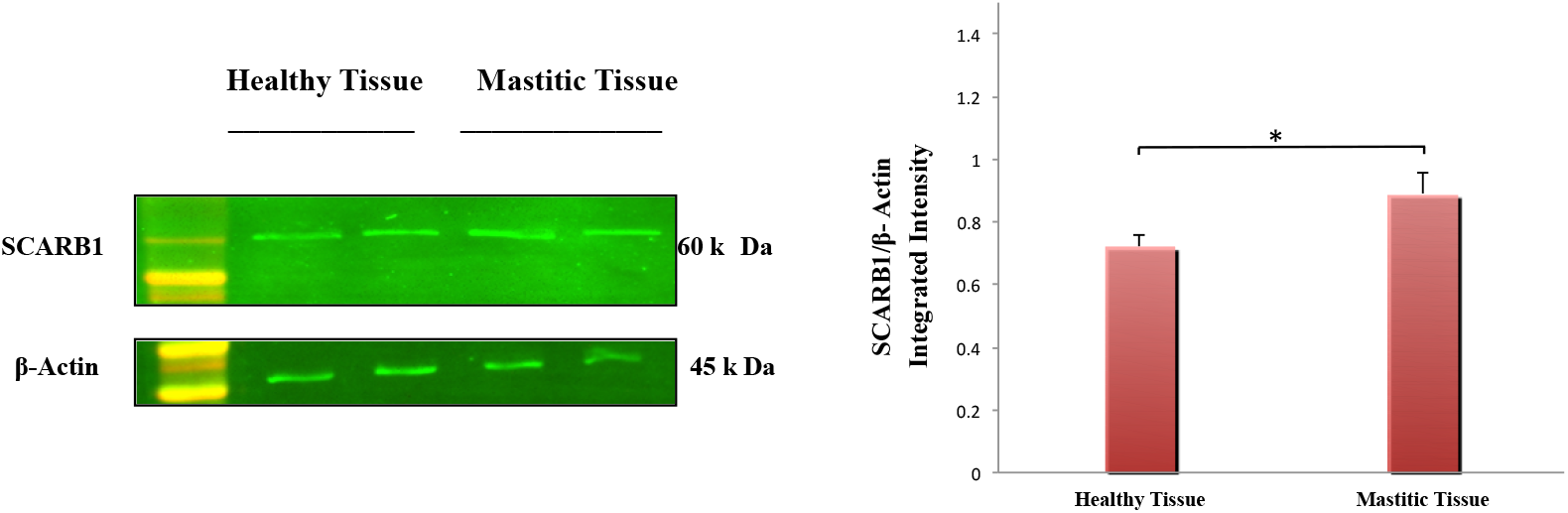
SCARB1 protein expressions were increased in mammary tissue obtained from mastitic goats as compared to healthy goat mammary tissues. The graph shows the densitometry quantification of SCARB1/β-actin, as the log2 fold change difference compared to the control. The data is presented as the mean ± SEM for duplicate samples. \**P* < 0.05.

#### In vitro SCARB1 expression in GMECs

The *in vivo* studies validated the involvement of the SCARB1 receptor in *E. coli*-induced coliform mastitis. In order to understand the role of SCARB1 during *in vitro E*.*coli* induced infection, the GMECs were stimulated at multiple time points and MOIs in this study. At MOIs of 1:50, 1:100, 1:300 and 1:500, *E*.*coli* did not induce cell necrosis or apoptosis while as at 1:1000 cytotoxicity was induced at 3 h, 6 h, and 24 h, which were determined by MTT cytotoxicity assay **(S1, Fig. 5)**. Following this, MOIs at 1:100, 1:300 and 1:500 was selected as the infection points and 3 h, 6 h (early time points) and 24 h (late time points) were included.

Stimulation of GMECs with hk *E*.*coli* significantly upregulated the mRNA expression of SCARB1 after 6 h as compared to non-stimulated cells. SCARB1 mRNA expression at different MOIs (1:100, 1:300 and 1:500) were significantly increased (1.114-3.689 folds, *P* < 0.05) by the stimulation at all the MOIs when compared to the non-infected control. Similarly, on infection with hk *E*.*coli* at 1:100, 1:300 and 1:500 MOIs after 6 h, protein levels of SCARB1 were significantly increased as compared to a non-infected control group with an increase in MOI (*P* < 0.05, *P* < 0.01) **(Fig. 6)**. However, on stimulation with hk *E*.*coli*, SCARB1 mRNA levels declined (0.65-0.115 folds, *P* < 0.05) after 24 h at all the MOIs when compared to the non-infected control cells. Similarly, SCARB1 protein levels were significantly decreased on infection with hk *E*.*coli* after 24 h as compared to the non-infected control group **(S2, Fig. 2b)**. No significant changes in SCARB1 mRNA and protein expression were notable between the non-infected control and hk *E*.*coli* group at 3 h post-infection (1.176-1.226 folds, *P <* 0.05) **(S2, Fig. 2a)**.

**Fig. 6.**
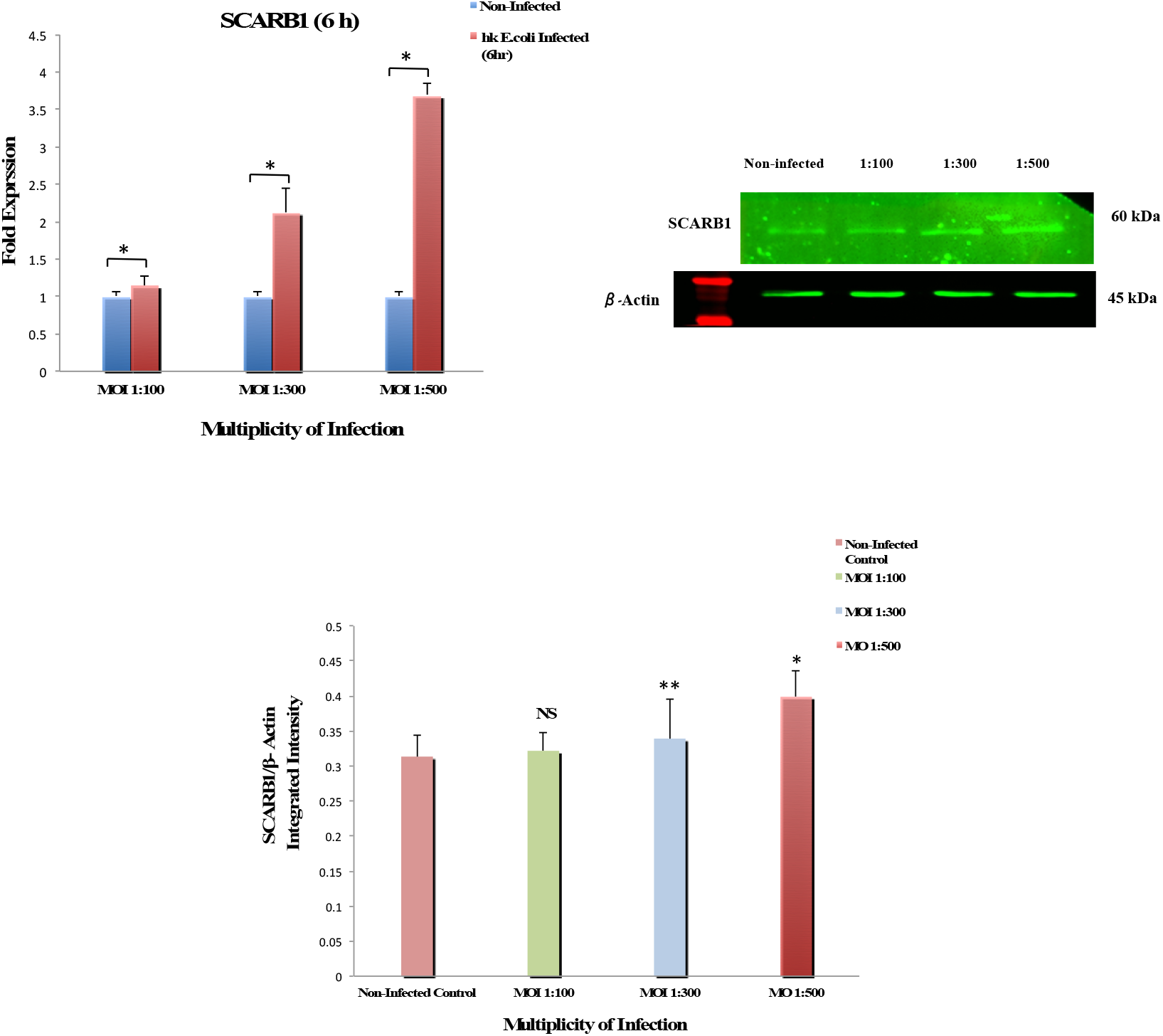
*E*.*coli* induced SCARB1 expression in GMECs. Cells treated with hk *E*.*coli* for 6 h at increasing MOIs (1:100, 1:300 and 1:500) show a significant increase in mRNA expression and protein levels of SCARB1. The graph shows the densitometry quantification of SCARB1/β-actin, as the log2 fold change difference compared to the non-infected control obtained from three separate blots. Quantitative PCR data were normalized to GAPDH and β-Actin. The values are the mean ± SEM for triplicate samples. \**P* < 0.05, \*\**P* < 0.01, and not significant (NS).

Additionally, LPS stimulation of GMECs significantly upregulated the expression of SCARB1 mRNA expression in a dose-dependent manner at different LPS concentrations (1, 10 and 50 μg/ml) at 6 h as compared to non-LPS treated cells (1.30-2.28, *P* < 0.05) **(S2, Fig. 2c)**. This indicates that 6 h time point is ideal for the significant expression of SCARB1 triggered by LPS of *E*.*coli* in GMECs.

### SCARB1 affects the response pattern of TLR4-MyD88 pathway

To understand if knockdown of SCARB1 affected the TLR4 pathways, the changes in SCARB1 mRNA expression and protein expression by knockdown of SCARB1 in GMECs at time points of 24, 48 and 72 h were observed **(S2, Fig. 3b)**. SCARB1 expression levels were significantly decreased after 72 h of transfection with esiRNA-SCARB1 (*P* < 0.05). NC-esiRNA-GFP expression was observed in GMECs treated with NC-esiRNA for 72 h (S2, Fig. 3a).

SCARB1 expression was detected at MOIs (1:100, 1:300 and 1:500) for 6 h before treating the cells with NC-esiRNA and esiRNA-SCARB1 at 72 h (*P* < 0.05) **(S2, Fig. 3c)**. The 1:500 MOI is the ideal MOI concentration to upregulate the expression of SCARB1 mRNA in the hk *E*.*coli* group.

Both the SCARB1 and TLR4 mRNA levels were influenced by the manipulation of SCARB1 expression on hk *E*.*coli*-induced GMECs for 6 h; however, no changes were noted in esi-SCARB1 cells without bacterial stimulation. In contrast to the hk *E*.*coli* -stimulated NC group, the SCARB1 and TLR4 mRNA levels (*P* < 0.05) declined dramatically in the NC group. In the deficiency groups both SCARB1 (*P* < 0.05) and TLR4 (*P* < 0.05) levels declined **(Fig. 7a,b)**. SCARB1 protein levels were also influenced by the manipulation of SCARB1 expression. The protein level in the hk *E*.*coli* - stimulated NC group was more as compared to the negative control (NC) group (*P* < 0.05). While as, the protein expression decreased in the esi-SCARB1 deficiency group (*P* < 0.01) **(Fig. 7a)**.

**Fig. 7.**
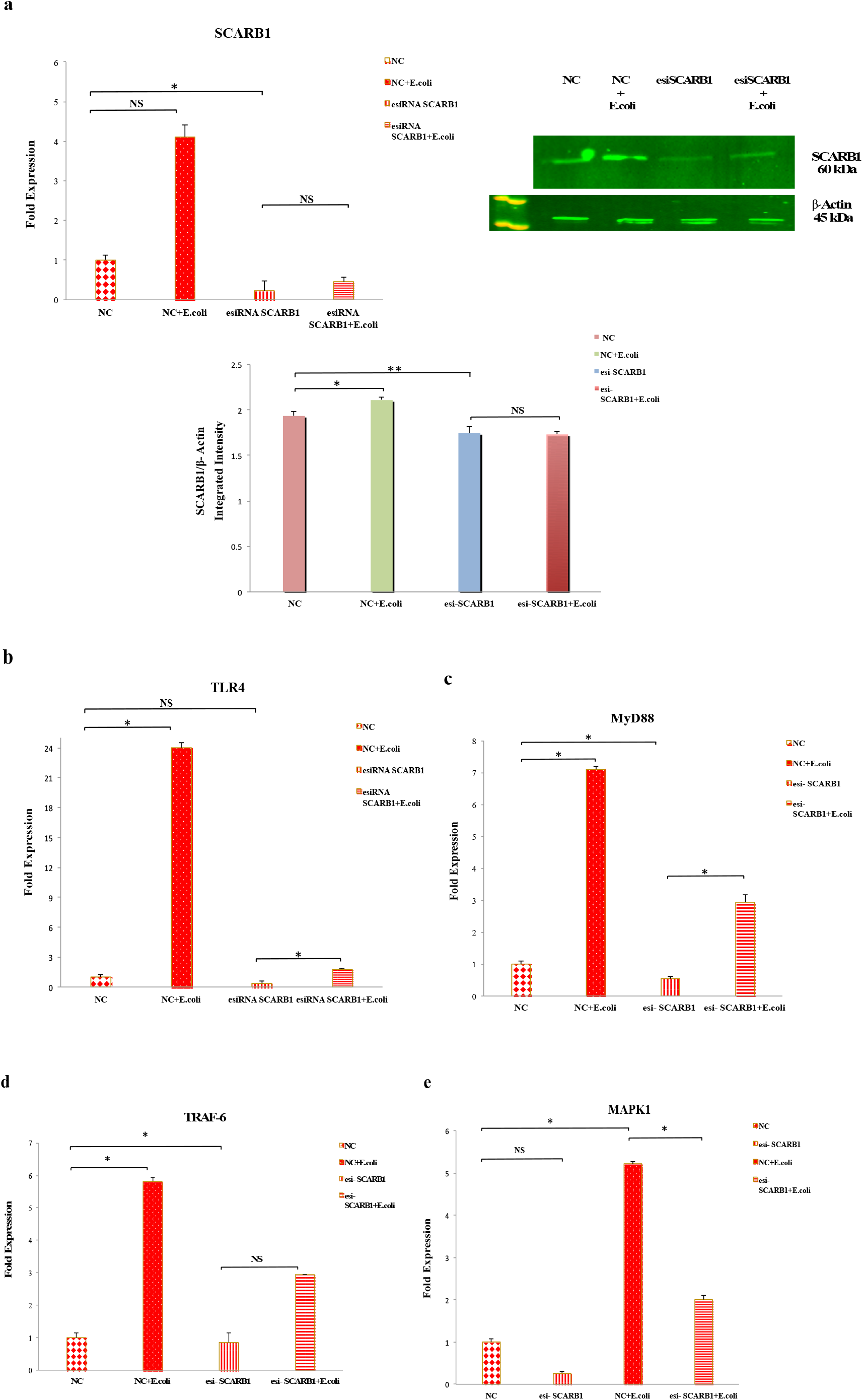
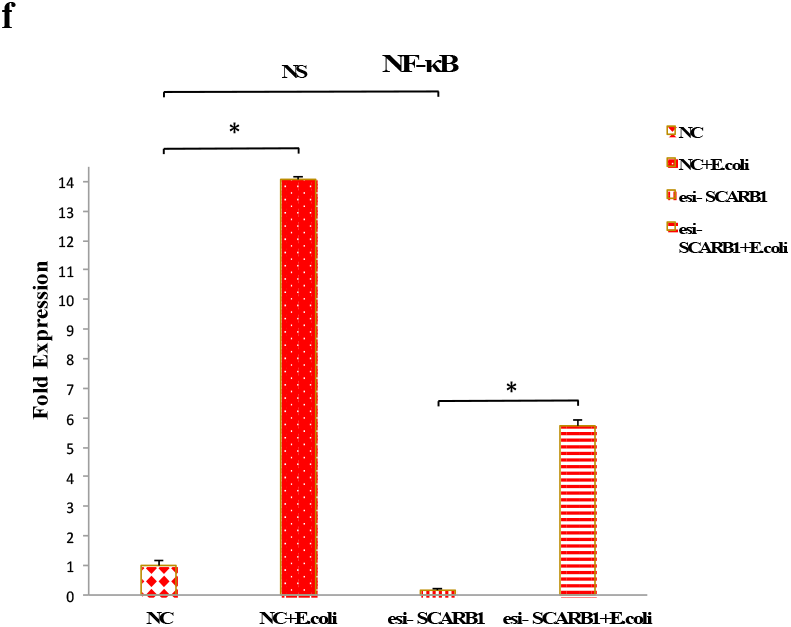
The silencing of SCARB1 regulates the MyD88 pathway via NF-κB and MAPK1 activation in GMECs exposed to *E*.*coli* infection. **a**, The significant changes in SCARB1 mRNA and protein levels following the silencing of SCARB1 in GMECs and infection with *E*.*coli* at 1:500 MOI. The graph shows the densitometry quantification of SCARB1/β-actin, as the log2 fold change difference obtained from three blots. b, TLR4 mRNA levels were influenced by knockdown of SCARB1 in GMECs, on infection with *E*.*coli* at 1:500 MOI. **(c, d, e, f)**. Expression of MyD88, TRAF6, MAPK1 and NF-κB were influenced by knockdown of SCARB1 in *E*.*coli* infected GMECs. Quantitative PCR data were normalized to GAPDH and β-Actin. All data are presented as the mean ± SEM from three experiments. \**P* < 0.05, \*\**P* < 0.01 and not significant (NS).

MyD88 and TRAF6 expression was increased in the hk *E*.*coli*-treated NC group compared with the NC group (*P* < 0.05) however; the expression was decreased in the esi-SCARB1 group compared to the esi-SCARB1+hk *E*.*coli* -treated groups (*P* < 0.05) **(Fig. 7c,d)**. Expression of `MAPK1` **(Fig. 7e)** `and NF-κB` **(Fig. 7f)** `is declined in deficiency groups as compared to control groups` (*P* < 0.05). These above-mentioned results indicate that knockdown of SCARB1 along with TLR4 affects MyD88 downstream signaling and that the MyD88, TRAF6, MAPK1 and NF-κB expression is stimulated only after *E*.*coli* -infection in GMECs.

### SCARB1 affects the response pattern of the TLR4-TRIF pathway

To evaluate the role of silencing of SCARB1 on the MyD88 independent pathway or TRIF pathway in the *E*.*coli* infected cells, relative expression of TRIF pathway genes was evaluated. The expression of TRIF and TRAF3 as compared with the NC group was increased in the hk *E*.*coli*-treated NC group (*P* < 0.05) and the expression was diminished in the esi-SCARB1 group compared to the esi-SCARB1+hk *E*.*coli* -treated groups (*P* < 0.05). In contrast to the hk *E*.*coli*-stimulated NC, the IRF3 mRNA levels declined dramatically in the deficiency groups (*P* < 0.05) **(Fig. 8)**. These results infer that the TRIF pathway genes and that of MyD88 pathway genes follow the same expression trend. Also, NF-κB is activated on TRIF pathway activation but in the late phase. These above-mentioned results indicate that knockdown of SCARB1 affects TRIF downstream signaling and that the TRIF, TRAF3 and IRF3 expression is stimulated only after *E*.*coli* -infection in GMECs.

**Fig. 8.**
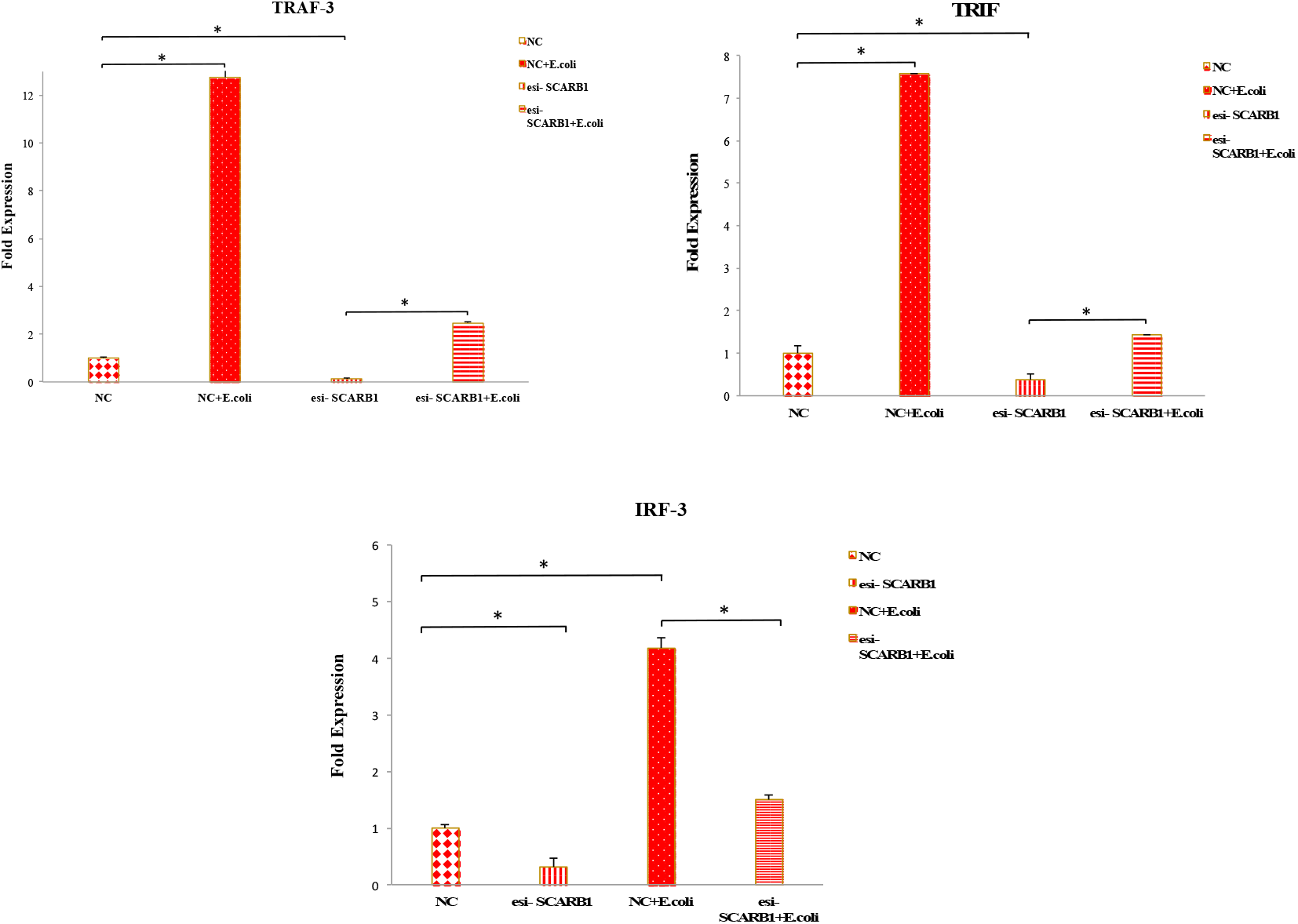
The silencing of SCARB1 regulates the TRIF pathway via IRF3 and NF-κB activation in GMECs exposed to *E*.*coli* infection. Knockdown of SCARB1 on infection with *E*.*coli* influences the expression pattern of TRAF3, TRIF and IRF3. Quantitative PCR data were normalized to GAPDH and β-Actin. All data are presented as the mean ± SEM from three experiments. \**P* < 0.05.

### SCARB1 regulates the expression of proinflammatory mediators on bacterial stimulation in GMECs

After depletion of SCARB1 expression in GMECs followed by 6 h stimulation with *E*.*coli*; the expression of proinflammatory mediators was assessed. SCARB1 manipulation affects the mRNA levels of cytokines such as interleukin 8 (IL-8), TNF-α and INF-β. The mRNA expression of all cytokines increased on infection with *E*.*coli* for 6 h as compared to the non-infected NC group (*P* < 0.05). IL8 and TNF-α are the major pro-inflammatory mediators produced on activation of TLR4-MyD88 pathway, while as on activation of TLR4-TRIF pathway INF-β is expressed. In the SCARB1 knockdown groups, the mRNA levels of the IL8 **(Fig. 9a)** was elevated after the cells were treated with *E*.*coli* for 6 h as compared to esi-SCARB1 cells without bacterial stimulation (*P* < 0.05). However, SCARB1 silencing has no significant effect on the mRNA expression of TNF-α and INF-β (*P* < 0.05) **(Fig. 9 b,c)**. This may be due to the fact that TNF-α and INF-β expression may depend on the activation of other infection pathways on *E*.*coli* stimulation in GMECs. These results indicate that SCARB1 regulates the production of proinflammatory mediators in GMECs in response to *E. coli*.

**Fig. 9.**
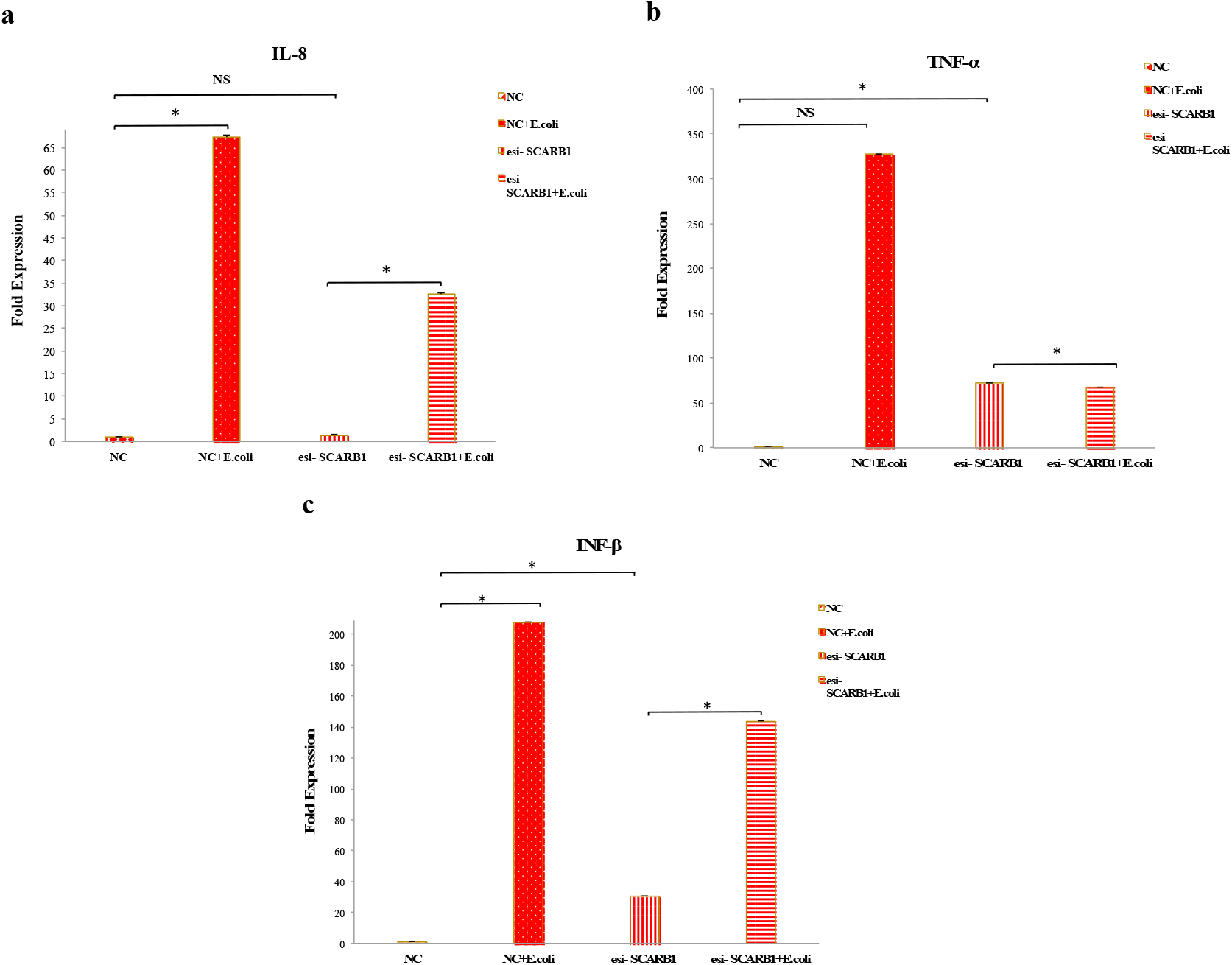
The silencing of SCARB1 followed by infection with *E*.*coli* in GMECs influences the expression of Proinflammatory mediators. The mRNA expression levels of the proinflammatory mediators **a**, IL8, **b**, TNF-α, **c**, INF-β in GMECs treated with *E*.*coli* at MOI 500 for 6 h prior to SCARB1 knockdown. Quantitative PCR data were normalized to GAPDH and β-Actin. All data are presented as the mean ± SEM from three experiments. \**P* < 0.05 and not `significant` (NS).

### Expression validation by RNA seq data

#### SCARB1 expression

The expression values of SCARB1 for control vs. *E*.*coli* infected samples were measured as fragments per kilobase of exon per million fragments mapped (FPKM) using Cufflinks v2.1.1 package with absolute log2 (fold change) of -0.050414 (49).

#### Protein-protein interaction analysis

In this study on *E*.*coli* induction of GMECs, TLR4, TRAF3 and TRAF6 are significantly overexpressed as compared to the non-induced cells and the PPI network also suggests the key role of TLR4 and TRAF6/3 in the *E. coli* infection. The SIGNOR database also suggests that *papain*-like proteinase and *NF-kB-p55/p50* signaling are involved in the process, which needs further evaluation **(S2, Fig. 4)**.

#### Pathway analysis

The KEGG enrichment analysis suggests *Toll-like receptor cascades* and *MAPK signaling pathway* **(S2, Fig. 5 and Table 3)** was significantly enriched. The terms enriched in Reactome and GO database are provided in **S2, Table 4 and 5** respectively, which also suggests the involvement of TLRs and MAPKs in *E*.*coli* Infection. Moreover, these critical pathways known for mounting an effective immune response may also aid in evading recognition and virulence and thus destabilize the host immune system by targeting signal transduction factors that need further validation (67).

### SCARB1 mediated *E. coli* internalization/endocytosis in GMECs

The SCARB1 mediated internalization of bacteria in GMECs was assessed by manipulating the expression of the SCARB1 gene i.e. treating cells with esi-SCARB1 followed by infection with live *E*.*coli*. In the esi-SCARB1 treated group, the number of colonies on LB agar plates was less in number as compared to mock NC-esiRNA treated cells **(Figure 10a)**. It is because SCARB1 might be involved in the internalization of bacteria. On the other hand, the deficiency of this receptor prevents entry of *E*.*coli* thus decrease in the rate of bacterial internalization in the presence of antibiotics.

**Fig. 10.**
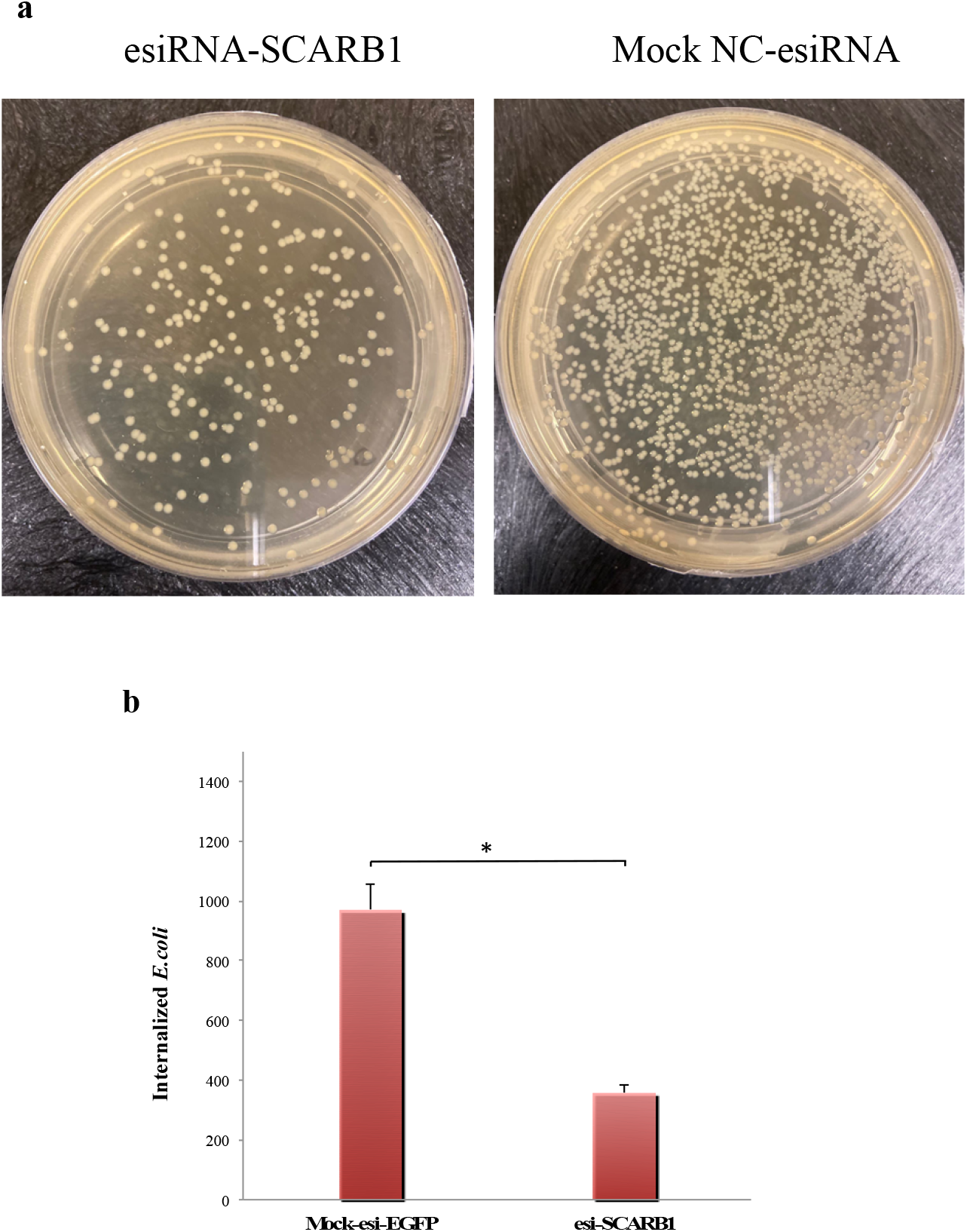
**a**, The GMECs were treated with NC-esiRNA mock and esi-RNA-SCARB1 followed by incubation with *E. coli* for 2 h. After cell lysis, the lysate (10^8^ cfu/ml) was plated on LB agar. **b**, The data is presented as the mean ± SEM from three experiments. \**P* < 0.05.

### Prediction of SCARB1 Structure

Using the BLAST program against the RCSB-PDB database we identified the five best templates for homology modelling (4Q4B, 4F7B, 4F7B, 4TW2 and 4TVZ). The sequence similarities between all five templates of SCARB1 were approx. 35-36%. Using Homology modelling and the Rosetta program the predicted structure show low coverage and low Global Model Quality Estimate (GMQE) of ∼61% **(S2, Fig. 6a-b)**. The structure predicted from Multi-threading (I-Tasser Program) covers the full length and the core region has a higher degree of confidence (C-Score of core region=1.5). Ramachandran validation shows 71.1% residues are in the core region, 27.8% residues are in the allowed region and 1.1% residue is in the disallowed region **(Fig. 11)**. The PDB file is provided in the supplementary AA.

**Fig. 11.**
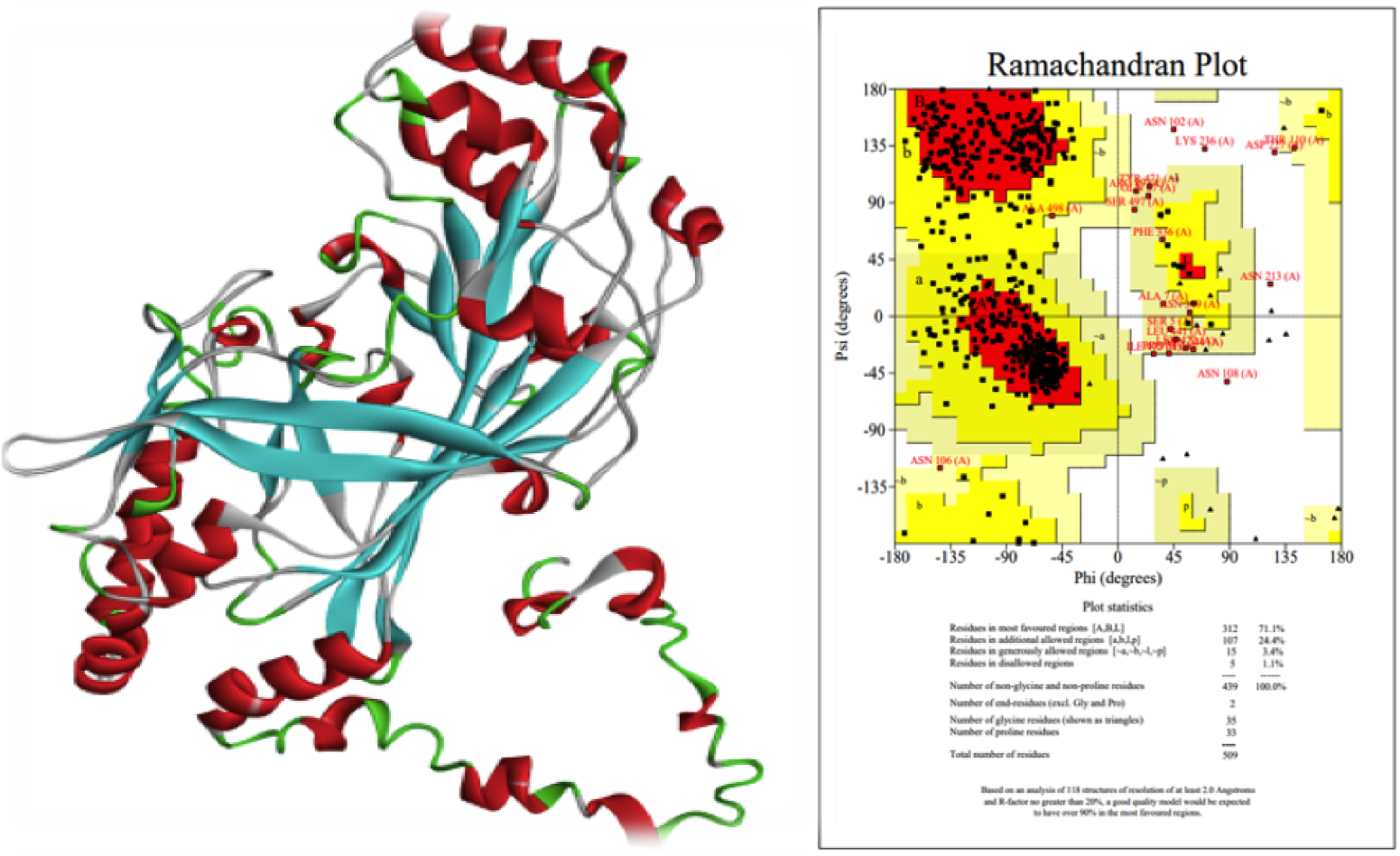
Structure predicted using I-Tasser Program. 71.1% residues are in the core region, 27.8% residues are in the allowed region and 1.1% residues are in the disallowed region.

## Methods

### Experimental animals

#### Bacterial identification

The Bakarwal goats with mammary gland infection were screened based on extreme clinical symptoms from Veterinary Clinical Complex (VCC), Faculty of Veterinary Sciences & Animal Husbandry (FV.Sc. & A.H), Sher-e-Kashmir University of Agricultural Sciences and Technology of Kashmir (SKUAST-K), Shuhama, Alustang.

Mastitic milk was collected aseptically using sterile vials and subjected to somatic cell counting (SCC). Coliform mastitis was determined through selective media (MacConkey agar and EMB agar) followed by gram staining and biochemical tests: Indole Methyl red Voges-Proskauer Citrate tests (IMViC) (38).

#### Tissue collection

For *in vivo* study, mammary gland tissue from Bakarwal goats (1.5-3 years, 25-65 kg), healthy control (n = 2) and mastitic (n = 2) in second parity, were selected for tissue collection. Mammary tissue biopsates were procured from the right or left mammary glands of healthy and mastitic goats through a biopsy procedure under sterile conditions (39) and were immediately frozen in liquid nitrogen for total protein extraction as already mentioned.

#### Milk collection

To harvest goat milk epithelial cells (GMECs), filtered raw goat milk was obtained from early lactating healthy Bakarwal goat from Mountain Research Centre for Sheep and Goat (MRCSG) of Sher-e-Kashmir University of Agricultural Sciences and Technology, Kashmir (SKUAST-K), Shuhama in sterile conditions. The teats were cleaned with 70% ethanol and iodine solution and the first few strips of milk were discarded before collection of the milk sample.

### Isolation and culture of Mammary Epithelial Cells from goat milk

About 50 ml of goat milk was collected in sterile falcon centrifuge tubes and kept on ice before processing. The protocol for the processing of milk has been accepted for a patent under Application No.201911013320 A, Dated: 09/10/2020. The cell isolation process was performed immediately after milk collection. Milk fat was separated by centrifugation and skimmed milk was removed. Pellets were pooled and suspended in sterile Dulbecco’s phosphate-buffered saline (DPBS; Himedia) followed by centrifugation. After washing, the cell pellet was suspended added to T25 flask (BD, Falcon) in Dulbecco’s modified Eagle medium (DMEM; Sigma-Aldrich) supplemented with fetal bovine serum (FBS) (Himedia), a wide range of antibiotics and growth factors in the water-jacketed incubator (Thermo Scientific) at 37°C and 5% CO2 humidity. After 24 h, washing was done twice with DPBS, and a fresh culture medium with all the antimicrobials and FBS was added to the culture flask. The culture medium was changed every 3 days and at 80% confluency, the goat mammary epithelial cells were detached by trypsinization using 1X trypsin-EDTA (Himedia). The MECs were passaged at a density of 5×10^5^ cells in culture flasks and continuously subcultured up to 15 passages. The cells were split in a ratio of 1:4 for routine culture. The cell line was cryopreserved in Liquid N_2_ in 1-1.5 ml aliquots in freezing media consisting of 70% DMEM, 20% FBS and 10% DMSO (Sigma-Aldrich).

### Growth, Proliferation and Morphological Characteristics of GMECs

The growth and proliferation pattern of GMECs was assessed by growth curve analysis as described previously (40). In brief, 1×10^4^ cells/well of early and late passage were seeded in 12-well culture plates (BD Falcon). Cell number and viability were examined each day in triplicate wells using trypan blue exclusion and cell proliferative assay (# Cat. no. T8154, Sigma-Aldrich). The assay was performed for 10 d post-seeding i.e., at 24, 48, 72, 96, 120, 144, 168, 196, 216 and 240 h. The average cell number calculated every 24 h was used to draw the growth curves over time.

For morphological and subcellular cell identification, early and late passage GMECs (P2, P3, P4, P13, P14 and P15) on the 7th day of culture were fixed with 4% formaldehyde solution and stained in 20% Giemsa solution (Sigma-Aldrich) for basic morphological identification.

The MTT [3-(4,5-dimethylthiazol-2-yl-2, 5-diphenyltetrazolium bromide)] assay was performed to check the viability and rate of proliferation for early passage, P2 and late passages GMECs, P13, P14 and P15 (41). Approximately, 1×10^5^ cells were seeded in 96 well plates. Subsequently, 10*μ*l MTT (50 mg/ml) (Sigma-Aldrich) was added to each well and kept at 37°C for 4 h. After 0.5 h MTT Solvent (10% SDS in 0.01M HCl) was added. The assay was performed for the next 7 days and each day absorbance was read at 570 nm using Cytation 5 Cell Imaging Multi-Mode Reader (Agilent, BioTek). The assay was performed in triplicate.

### Immunofluorescence of Cytokeratin 18 (CKT-18)

GMECs, fibroblasts (Negative control) and MCF 10A (Positive control) cells were cultured in 12 well culture plates at a density of 2×10^4^ cells/well and grown at 37°C in a humidified CO_2_ incubator until 60-70% confluency. Goat Fibroblasts were obtained from *ATCC* (Ch 1 Es (NBL-8) and MCF 10A cells were provided by Dr L.S Shashidhara, ICAR, Pune, India). IF was performed for detection of epithelial cell-specific marker CKT-18 with specific antibodies (42). GMECs of early passage P2, P3 and late passage P15, fibroblasts and MCF 10A were washed with DPBS (Himedia), before fixation with 4% formaldehyde at room temperature. For detection of CKT-18 fixed cells were permeabilized with 0.1% Triton X-100 (Himedia) at room temperature. Cells were washed thrice with 0.05% TBS (Tween-20 in DPBS) (Himedia). After washing, cells were blocked with 5% FBS and 2% bovine serum albumin (BSA) (Sigma-Aldrich) for 1 h. GMECs and fibroblasts were incubated overnight at 4°C with cytokeratin-18 monoclonal antibody (# Cat. No. MA1-19031, Thermo Scientific, USA). MCF 10A cells were incubated with CKT-18 Monoclonal Antibody (# Cat. No. RCK106, Thermo Scientific, USA). The next day, wells were washed thrice with TBS. GMECs, fibroblasts and MCF 10A cells were incubated with the FITC bound anti-mouse IgG secondary immunoglobulin (Fc specific) (Cat # F4143, Sigma-Aldrich) secondary antibody for 2 h at room temperature away from light. Thereafter, nuclei were counterstained with 4′, 6-diamidino-2-phenylindole (DAPI) (Sigma-Aldrich) in 1% block solution and incubated in dark at room temperature. Images were acquired using the FLoid cell Imaging workstation (Thermo Scientific) at 100x.

### Chromosomal Analysis

Karyotype analysis was performed to determine the chromosome number throughout the passages. It was performed at the Department of Advanced Centre for Human Genetics SKIMS Kashmir described previously (43). Growing GMECs at early (P3) and late passages (P12) were treated with colchicine (10 μg/ml) (Sigma-Aldrich) for about 7 h. Then the cells were trypsinized and treated with warm hypotonic KCL solution (56%) at 37°C followed by centrifugation. Cells were fixed on a glass slide and the sample slides were Giemsa stained. The slides were visualized on a phase-contrast microscope for the chromosomes and karyotyping analysis through cytovision genus software (Applied Imaging, USA).

### Inducing differentiation of GMECs

Goat mammary epithelial cells at passage 2 were induced to differentiation towards milk-secreting cells (44). In addition to normal constituents of growth media, induction media was supplemented with the combination of 5μg/ml prolactin (Sigma-Aldrich), 5μg/ mL insulin (Himedia), 5μg/mL hydrocortisone and 20 ng/ml EGF (Sigma-Aldrich) to induce the expression of CSN-2. Culturing the cells in induction media for 5 days induced differentiation. After 5 days the RNA was isolated from cells, cDNA was synthesised and gene expression analysis of CSN-2 was done by RT-qPCR. Prior to RNA isolation, the differentiated cells were visualized under an Inverted Microscope (IX73, Olympus, Japan).

### Transfection of the vector containing enhanced green fluorescent protein (EGFP) in GMECs

Cells were seeded in 12 well culture plates (BD, Falcon cat No 353043) 24 h before transfection by Turbofect (#Cat. No. R0533, Thermo Scientific,). The 4.7 kb vector (EGFPC1) used in the current study has EGFP under CMV promoter (Courtesy: Dr Khalid Masoodi’s K-Lab, SKUAST-K, Shalimar). The vector was upscaled in the *E. coli* (DH5 alpha) strain. After 24 h, the plasmid DNA was added with transfection reagent and serum-free DMEM (Sigma-Aldrich,) to cells for 24 h by strictly following manufacturers’ instructions. Thereafter cells were left undisturbed in humidified CO_2_ incubator at 37°C and 5% CO2. The cells were observed 24 h after transfection under an Inverted Microscope (IX73, Olympus, Japan) (42).

### Cell Culture and Bacterial Challenge

*E. coli* challenged GMEC model was used to simulate the *in vitro* experimentation. *E*.*coli* (ATCC® 25922™) were cultured to an optical density (OD_600nm_ ≈ 0.6) in Luria-Bertani (LB) medium at 37° C with constant shaking at 200 rpm. Bacterial cells were collected by centrifugation at 1500 rpm for 5 min, washed 2 times DPBS, and then resuspended in sterilized DPBS followed by serial dilutions (10^1^ - 10^8^). At passage 2, the cells were seeded with supplemented DMEM media in 12 well plates (BD Falcon) at a density of 8×10^4^ cells/cm^2^ at 37°C in an incubator for *E. coli* infection. *E. coli* was heat-killed (hk) at 63°C for 30 minutes before infection to avoid overgrowth during the period of infection. Cells were challenged with hk bacteria following a previous study (45). At 80% confluence, the growth medium was replaced by infection medium (same as the growth medium but without antibiotics) and cells were treated with bacteria (10^6^ cfu/ml) at thrqee time points (3 h, 6 h, 24 h) and at multiplicities of infection (MOIs) of 1:100, 1:300 and 1:500. Cytotoxicity assay (MTT) was carried out for MOI selection. Cells were treated with LPS O55:B5 (Sigma-Aldrich) at different concentrations (1, 10, 50 μg/ml) for 6 h (26). Uninfected control (infected with the same volume of heat-inactivated infection media but without bacteria) was also included. Experiments were performed in triplicate.

### Confirmation of the Bacterial infection

#### Adhesion assay

A bacterial Adhesion assay was performed to confirm adherence by visualization of the adhered bacteria on infected GMECs (46). About 1.5 × 10^5^ GMECs were seeded in 12 well culture plates (BD Falcon). At 80% confluency, the growth medium was replaced by an infection medium (Medium with bacteria). Monolayers/wells were infected with hk bacteria (10^6^ cfu/ml) at an MOI of 1:100 for 3 h at 37°C with 5% CO_2_. Uninfected control wells with infection media with no bacteria were also included. The wells were stained with 20% Giemsa stain (Sigma-Aldrich) for 30 minutes and visualized under Inverted Microscope (IX73, Olympus, Japan) at 100x.

#### Cellular response to bacterial challenge

Prior to investigating the immune response of GMECs to infection, we wanted to confirm if our bacterial challenge influenced the expression of immune-responsive genes in these cells. Real-time expression of TNF-a and IL8 mRNA was assessed through RT-qPCR following challenge by hk *E. coli* for 3 h at 1:100 MOI. The Primers details are given in **(S1, Table 2)**. The rest of the calculations for differential quantitation of genes is the same as mentioned.

### SCARB1 knockdown and cell treatment

For SCARB1 knockdown, MISSION esiRNA (esi-SCARB1) was designed and manufactured by Sigma-Aldrich (47). Cells were treated with EGFP tagged esiRNA that served as an NC (NC-esiRNA-EGFP, Sigma-Aldrich). About, 2.5 × 10^5^ primary GMECs were seeded in 6 well plates (BD, Falcon cat No 353043) to 70% confluence and transfected with (2μg) esi-SCARB1 and NC-esiRNA-EGFP using X-tremeGENE 360 Transfection Reagent (Sigma-Aldrich) according to the manufacturer’s instructions. Opti-MEM I Reduced Serum Medium (Thermo Scientific) was used for the formation of a transfection complex. The esiRNA sequences, as well as transfection reagents used, are provided in **(S1, Table 1)**. The time-dependent knockdown efficiency was detected after 24, 48 and 72h in cell culture by RT-qPCR and western blotting as described. SCARB1 expression levels were significantly decreased after 72 h of transfection with esiRNA-SCARB1. NC-esiRNA-EGFP expression was observed in GMECs treated with NC-esiRNA for 72 h. Then, for ideal MOI selection, in a subsequent experiment, after 72 h of transfection with NC-esiRNA-EGFP or esi-SCARB1, GMECs were treated with hk *E*.*coli* for 6h at increasing MOIs (1:100, 1:300 and 1:500). NC-esiRNA-EGFP expression was observed in GMECs treated with NC-esiRNA for 72h under an Inverted fluorescent Microscope (IX73, Olympus, Japan). Experiments were performed in triplicate.

### Western blot

Whole protein from mammary tissue and GMECs was extracted with NP-40 (Himedia) lysis buffer supplemented with PMSF (Himedia), PIC (Sigma-Aldrich) and NaF (Himedia). Protein concentrations were quantified in Qubit 2.0 using a Protein Assay Kit (Thermo Scientific) according to the manufacturer’s instructions. Western blotting was performed using rabbit monoclonal Anti-Scavenging Receptor SR-BI antibody (# Cat. no. ab52629, Abcam) and beta Actin mouse Monoclonal Antibody (AC-15) (# Cat. No. MA1-91399, Thermo Scientific). The secondary antibodies used were Anti-rabbit IgG (H+L) (DyLight^™^ 800 4X PEG Conjugate) #5151 and Anti-mouse IgG (H+L) (DyLight^™^ 800 4X PEG Conjugate) #5257 (Cell Signaling Technology) respectively. All the antibodies were used according to the manufacturer’s recommendations. Protein bands were visualized using ChemiDoc MP (BioRad) and intensity was evaluated by Image-Lab software 6.0. Experiments were performed in triplicate.

### Total RNA extraction and cDNA synthesis

Total RNA from GMECs was extracted using the RNeasy Micro kit (# Cat. No. 74004, Qiagen) according to the manufacturer’s instructions. RNA was quantified on Nanodrop Lite (Thermo Scientific) and before cDNA synthesis. cDNA was synthesized with an equal concentration of RNA (1.5 μg/μl) in all the samples using Thermo Scientific RevertAid First Strand cDNA Synthesis Kit (Lithuania) by oligo dT primers according to the manufacturer’s instructions. cDNA synthesis was validated through conventional PCR in a thermocycler (Applied Biosystems). PCR was performed with GoTaq® Green Master Mix (2X) (Promega Corporation) using gene-specific primers **(S1, Table 2)** and cDNA as templates.

### RNA Sequencing (RNA-Seq)

To further affirm the role of SCARB1 and TLRs in infection we used RNA-Seq data from accession No. GSE167591. We utilized one control sample (Non-Infected - Primary Mammary Epithelial Cells) and three *E*.*coli* infected Primary Mammary Epithelial samples. RNA extraction was carried out with Trizol Reagent (Ambion, USA) according to the manufacturers’ instructions, which was followed by measuring the absorbance of RNA samples at 260 nm and 280 nm with a spectrophotometer (Thermo Scientific). Moreover, the quality and integrity were measured on the Agilent 2100 Bioanalyzer (Agilent, USA). The RNA integrity number (RIN) value of the samples used for library preparation was ≥8 and cDNA libraries were prepared using IlluminaTruSeq Stranded mRNA Sample Prep kit (Illumina, USA) from 4μg of total RNA according to the manufacturers’ instructions. The sequencing was performed at SciGenom Lab (Cochin, India) using Illumina HiSeq 2500. RNA-Seq data were filtered and analyzed as per the pipeline described previously (48, 49).

### Quantitative real-time PCR

Real-time quantitative RT-qPCR was performed using SYBR Green PCR Master Mix (KAPA™ SYBR® qPCR Kit, KapaBiosystems, Woburn, MA) according to the manufacturer instructions. The primers used for expression were already reported (*GAPDH, β-Actin, CSN-2, TLR4, TRIF, MyD88, IRF3, TRAF3, MAPK1, TNF-α, IL-8, NF-kB, TRAF6* and *INF-β)* (28, 50, 51, 52, 53, 54). *SCARB1* primers were designed using Primer3 Plus software. The SCARB1 PCR product was sequenced at AgriGenome Labs, Kerala, India and was subjected to BLAST on NCBI and was 100% similar to the reference sequence. The primers were purchased from Sigma-Aldrich. The primer sequence and annealing temperatures are given in **(S1, Table 2)**. All the qPCR reactions were performed in a 96-well plate on Lightcycler 480II (Roche™, Germany) and data normalized to GAPDH and β-Actin, which, were used as internal controls. Experiments were performed in triplicate.

### Statistical Analysis

The qPCR data were analyzed relative to the control using 2^−ΔΔCt^ method where Ct is the cycle threshold (55) and results are expressed as mean ± standard error of the mean (SEM). All of the results were achieved from at least three independent experiments. The data obtained in the current study were analyzed using a computer-aided statistical software package SPSS 20.0. The differences between means were analyzed by unpaired Student’s T-test and by one-way ANOVA. Statistical significance was declared at *P* < 0.05.

### Antibiotic protection assay

The ability of GMECs to internalize bacteria was detected under different SCARB1 expression conditions (mock and esi-SCARB1) (28). Before the GMECs were treated with the bacteria, the SCARB1 knockdown was carried out. Prior to the assay, the cells were washed with DPBS. Cells were seeded in a 6 well plate and cultured cells were incubated in DMEM containing 10% FBS, antibiotics and live *E. coli* (approximately, 10 bacteria/cell) in culture wells. After the cells were incubated for 2 h at 37 °C, the plates were washed with ice-cold DPBS (at least three times) and lysed in ice-cold water for 40 min on ice. Lysates were plated on LB agar dishes at various dilutions (10^2^–10^8^ cfu/ml) and incubated overnight at 37 °C. Experiments were performed in triplicate.

### Network Analysis

Four diverse databases were used for network analysis; protein-protein interaction (PPI) network was developed using STRINGDB (56). Pathway analysis and signaling networks were developed using the SIGNOR database (57). KEGG database (58) was used for the identification of pathways regulated by gene-set. To gain insight into the functional and enrichment analysis g: profiler webserver was used (59) using *Capra hircus* as reference.

### Structure prediction

Three different strategies of structure building were tested:

#### Homology Modeling (HM) or comparative modeling

HM is based on the principle that if two sequences share a high degree of similarity/identity, their respective structures are also similar. In this technique, we identified a proper template using BLAST program V2.12.0 (60), followed by protein sequence alignment, alignment correction/refinement, and structure prediction (backbone generation, loop modeling and side-chain modeling).

#### Iterative threading method

In this method, the structure templates were identified from RCSB-PDB (63) using a multi-threading approach by iterative template-based fragment assembly simulations. This method is implemented in I-TASSER (Iterative Threading ASSEmbly Refinement) (64).

#### Rosetta program

In this strategy, we utilized the Rosetta program (65) and the two-way approach for structure prediction. At first, the program uses a rigid body re-assembly approach, and the remaining parts of the structure were built using the *ab-initio* approach.

The predicted structures were refined by, correcting the bond order, creating di-sulphide bonds, filling missing side chains and loops and optimized using the energy minimization technique *OPLS3e* method. The structure was run through RAMACHANDRAN Plot Server (61) and PROCHECK (62) to test the quality of the predicted structure.

## Discussion

SCARB1 primarily as a lipoprotein receptor has been majorly known for scavenging lipoproteins (3, 4, 6, 7). Besides that, its role in attachment and internalization of a wide range of gram-negative, gram-positive bacteria and viruses on a variety of cell lines has been well established (15, 81, 82, 18) and its role in regulating phagocytic killing of gram-negative bacteria have also been reported (10). The *E. coli* mammary gland infections trigger an inflammatory response to the bacteria, which links exogenous pathogen with the endogenous immune system activation. Efforts at controlling mammary gland infections have focused on vaccine development, antibiotic treatment and infection management and at understanding the innate immune response on host-pathogen interactions (68, 69).

MECs are the first to confront the pathogen on entering the mammary gland (22, 23). In this study, milk-derived GMEC culture as an infection model was challenged with whole hk *E. coli*, unlike most studies where LPS is used as a virulence factor (83). The role of whole hk bacteria in innate immune response via Toll-like receptors has been shown (84) and heat inactivation seems to induce an elevated immune response (85) as well. In our study, we demonstrate that SCARB1 is involved in *E*.*coli* internalization/endocytosis and activation of proinflammatory response via TLR4 receptor in Goat mammary epithelial cells. On SCARB1 silencing, the rate of bacterial internalization decreased which corresponds to the less intracellular bacterial count in GMECs in the presence of antibiotics. Moreover, the response pattern of TLR4 pathways: MyD88 and TRIF pathways as well as the internalization of *E*.*coli* are inhibited by knockdown of SCARB1 expression. Thus, SCARB1 plays an important role in mammary gland innate immunity and might signify a therapeutic target to check mammary gland infections.

The PMECs (primary mammary epithelial cells) resemble the physiological environment of the *in vivo* cells more precisely and as a model, it is helpful in explaining molecular pathways associated with host-pathogen interactions (70). In this study, we developed an in house protocol to isolate and establish the pure PMEC culture from goat milk through a non-invasive method unlike the invasive methods of isolating cells from mammary tissue, which involves surgical intervention (71). There are only a few studies that attempted to isolate and culture MECs (72, 73, 74). Our method has been accepted for Patent under Application No. 201911013320 A for novelty in the process of culturing or propagating these cells in a monolayer manner which results in enhanced viability till a higher passage number i.e. 15, which is significantly high for a primary mammary epithelial cell culture. Thorough characterization of early and late passage cells present distinctive epithelial-like morphology, growth and proliferative characteristics, epithelial marker CKT-18, a non-transformed lineage throughout the passages i.e. 30-chromosome number (specific for *Capra hircus*) and differentiation into milk-secreting cells on lactogenic induction (75, 76, 77, 78, 79, 70). We also demonstrated that GFP could be efficiently transfected in GMECs, which can ease the physiological, and functional study of MECs through genetic manipulation *in vitro* (80). The established cell line represents a model useful for basic and applied research in mammary gland biology and biotechnology.

The g: profiler webserver was used using *Capra hircus* as a reference to gain insight into the functional and enrichment analysis. Here, we report new terms, which involve *papain-like proteinase* and *NF-kB-p55/p50* signaling to be involved in infection. *Papain-like proteinase* is one of two SARS-CoV-2 proteases, which plays role in the disruption of host response during infection and thus a potential target for antivirals (86). However, its function in *E*.*coli* infection needs to be further validated. The KEGG enrichment analysis, Reactome and GO analysis various Toll-like receptor-signaling cascades, TRAF6 mediated induction of NF-κB and MAP kinases (MAPK cascades) involvement in *E*.*coli* infection. Similarly, Protein-protein interactions suggest interactions of TLR4 pathway proteins on *E*.*coli* induction of GMECs such as TRAF3, TRAF6, TICAM1 (TRIF), MyD88 and MAPK1, which is consistent with the qPCR, based significant overexpression of these in infected condition vs. non-infected condition. Three unique protein structures of SCARB1 of *Capra hircus* were predicted through Homology modeling, Rosetta program and I-Tassser that will become the basis for future structural bioinformatics and docking studies.

In this study, on esi-SCARB1 pretreated cells (72 h) infection with *E*.*coli* at 1:500 MOI for 6 h could evoke an appropriate inflammatory response for a significant increase in SCARB1 and TLR4 mRNA levels and in activating the downstream factors NF-κB, MAPK1, IRF3 and proinflammatory cytokines and type I interferon. Cells challenged for 6 h with increasing MOIs of *E*.*coli* showed a significant increase in mRNA and protein expression of SCARB1. Similarly, cells on LPS treatment with increasing concentrations, also showed a significant increase in expression of SCARB1. However, cells challenged for 3 h with the same infection points showed no significant change in SCARB1 mRNA and protein expression. Also, at 24 h post-infection, SCARB1 mRNA and protein expression was significantly decreased on increasing MOIs, which is consistent with our RNA Seq data.

Only recently, a new function for SCARB1 has emerged as the inducer of cell signaling pathways upon ligand binding so no direct data regarding the role of *E*.*coli* binding to SCARB1 in TLR4 signal transduction has been reported. However, *in vitro* study has shown that SCARB1/SR-BI/CLA-1 plays an important role in *E*.*coli* and LPS induced proinflammatory signaling which is characterized by greatly elevated IL-8 secretion (8). Later *in vivo* studies reported that SR-BI contributes to the proinflammatory response by LPS in mice models and that the ability of SR-BI to recognize LPS contributes to TLR4 mediated pro-inflammatory response (87). Overexpression of SR-BI causes LPS to be transported to the *trans*-Golgi network (14) and SR-BI is associated with caveolin rafts, both are sites of TL4 localization. LPS internalization into the Golgi is associated with rapid MyD88 recruitment and LPS-TLR4 co-localization and is critically required for TLR4-NF-κB activation in intestinal epithelial cells (88). Also, blocking LPS binding to CLA-1 prevents pro-inflammatory cytokine responses (14). In this study, we report silencing of SCARB1 expression influence the expression of TLR4-MyD88 with subsequent production of pro-inflammatory molecules in GMECs triggered by *E*.*coli*.

TRAF6 is an important signaling molecule that transduces TLR4 signals to the NF-κB and MAPK pathways to directly modulate key cellular processes (89). TRAF6 has been reported to be associated with bifurcating the signal to the NF-κB and the MAPK activation pathways in LPS-stimulated mast cells (90). The relationship between SCARB1 and MAPK1 in *E. coli*-induced mastitis is an interesting finding. This article demonstrated that MAPK1 (ERK2) was influenced by the manipulation of SCARB1 expression during *E*.*coli*-induced activation in GMECs. It is shown that ERK1/2 potentially contributes to the enhanced inflammatory response of LPS-challenged (hSR-B tgn) transgenic mice (87). Blocking of ERK1/2 and NF-κB pathways prevent *E*.*coli* induced IL-8 production (91, 92) and TLRs present on the surfaces of intestinal epithelial cells appear to mediate LPS stimulation of ERK1/2 (93). Also, on stimulation of BMECs, TLR4 signalling cascade is activated which involves MyD88 and TRAF-6 that activates NF-κB and MAPK pathways (92). Moreover, mastitis studies in mouse models show activation of NF-κB and MAPK signaling pathways (95, 96). The downstream nucleic transcription factor NF-κB plays important role in exogenous-induced inflammation. In an *in vitro* study, *E. coli* and LPS strongly activated NF-κB followed by activation of proinflammatory cytokines (*e*.*g*. IL-8, and TNF-α) in BMECs (97, 76). This is consistent with our results where the cytokine TNF-α and chemokine IL-8 were regulated by SCARB1 in *E*.*coli*-induced inflammation in GMECs. In this current study, SCARB1 knockdown influenced the expression of TLR4, MyD88, TRAF6, MAPK1 and NF-κB in *E*.*coli*-stimulated GMECs, which confirmed that SCARB1 is involved in the TLR4-MyD88-dependent signalling pathway.

TRIF is responsible for mediating TLR4-dependent activation of several transcription factors, NF-κB and IRF3 leading to the production of cytokines and type-I IFN (98, 99) in response to LPS of *E*.*coli* (100, 101). This is consistent with our study where the expression of INF-β is regulated by SCARB1 in *E*.*coli*-induced inflammation in GMECs. On MAC-T cells on exposure to LPS, IRF3 is upregulated (27). While IRF3 activation is responsible for the expression of IFN-β most likely via the TRIF pathway on BMECs after LPS stimulation (28). Similarly, LPS treatment up-regulates IFIT3 (interferon regulatory factor 3) and upregulation of IRF3 with LPS and LTA (staphylococcal lipoteichoic acid) together (102). On the other hand, IRF3 activation by TLR4 is strictly dependent on the recruitment to TRIF to the adaptor TRAF3 that is well established (103). In our study, SCARB1 knockdown influenced the expression of TRIF, TRAF3 and IRF3 in *E*.*coli*-stimulated GMECs, which confirmed that SCARB1 is involved in the TRIF-dependent signalling pathway as well.

Only after TLR4 internalization, TRIF pathway is activated (36). On activation, TLR4 undergoes dynamin-dependent endocytosis where it triggers TRAM-TRIF–dependent signaling for IFN-β induction from an endosomal compartment. TRAF3 has been reported to localize in pleiomorphic cytosolic structures i.e. endosomes and is recruited to TRIF for IRF3 activation from endosomal location (37). So, here we can associate internalization/ endocytosis of bacteria with the TLR4-TRIF pathway activation, which on the other hand may suggest that SCARB1 in cooperation with TLR4-TRIF branch is involved in *E*.*coli* endocytosis in GMECs.

In conclusion, our results show that *E*.*coli* can exploit SCARB1 function to promote its cellular entry and SCARB1 regulates the *E*.*coli*-induced inflammation via the TLR4 “bipartite” receptor-mediated MyD88 and TRIF signaling pathways in GMECs. Thus, highlighting SCARB1 as a player in host defence and its potential role in antibacterial approaches to curb mammary gland infection.

## Acknowledgements

We are highly thankful to the Hon’ble Vice Chancellor Dr. Nazir Ahmad Ganai, SKUAST-K Shalimar, the Genetics Centre, SKIMS, Soura for technical support.

## Author contributions

Q.T., P.T.M., and S.M., performed experiments and wrote the manuscript. S.M.A., E.H., S.S., N.S., reviewed and edited the manuscript. B.B. analyzed the data. A.M., Z.A.K., M.H.Z., A.A.M., N.A.G., R.A.S. provided technical assistance. All authors approved the final version of the manuscript.

## Ethics approval

This study was performed according to the Institutional Animal Ethics Committee (IAEC), SKUAST-Kashmir, India.

## Data and materials availability

All data needed to evaluate the conclusions in the paper are present in the paper and/or the Supplementary materials. Additional data related to this paper may be requested from the authors

